# VLP-mediated delivery of structure-selected neoantigens demonstrates immunogenicity and antitumoral activity in mice

**DOI:** 10.1101/2023.09.07.556652

**Authors:** Ana Barajas, Pep Amengual-Rigo, Anna Pons-Grífols, Raquel Ortiz, Oriol Gracia Carmona, Victor Urrea, Nuria de la Iglesia, Juan Blanco-Heredia, Carla Anjos-Souza, Ismael Varela, Benjamin Trinité, Ferran Tarrés-Freixas, Carla Rovirosa, Rosalba Lepore, Miguel Vázquez, Leticia de Mattos-Arruda, Alfonso Valencia, Bonaventura Clotet, Carmen Aguilar-Gurrieri, Victor Guallar, Jorge Carrillo, Julià Blanco

## Abstract

**Background:** Neoantigens are patient- and tumor-specific peptides that arise from somatic mutations. They stand as promising targets for personalized therapeutic cancer vaccines. The identification process for neoantigens has evolved with the use of next-generation sequencing technologies and bioinformatic tools in tumor genomics. However, *in silico* strategies for selecting immunogenic neoantigens still have very low accuracy rates, since they mainly focus on predicting peptide binding to Major Histocompatibility Complex (MHC) molecules, which is key but not the sole determinant for immunogenicity.

**Methods:** We developed a novel neoantigen selection pipeline based on existing software combined with a novel prediction method, the Neoantigen Optimization Algorithm (NOAH), which takes into account structural features of the peptide/MHC-I interaction in its prediction strategy. Moreover, to maximize neoantigens’ therapeutic potential, neoantigen-based vaccines should be manufactured in an optimal delivery platform that elicits robust *de novo* immune responses and bypasses central and peripheral tolerance.

**Results:** We generated a highly immunogenic vaccine platform based on engineered HIV-1 Gag-based Virus-Like Particles (VLPs) expressing a high copy number of each *in silico* selected neoantigen. We tested different neoantigen-loaded VLPs (neoVLPs) in a B16-F10 melanoma mouse model to evaluate their capability to generate new immunogenic specificities. NeoVLPs were used in *in vivo* immunogenicity and tumor challenge experiments.

**Conclusions:** NeoVLPs can promote the generation of *de novo* antitumor-specific immune responses, resulting in a delay in tumor growth. Vaccination with the neoVLP platform is a robust alternative to current therapeutic vaccine approaches and a promising candidate for future personalized immunotherapy.

**WHAT IS ALREADY KNOWN ON THIS TOPIC:** Identification of highly immunogenic neoantigens is still challenging, currently available pipelines base their prediction on MHC-I binding affinity. Moreover, neoantigen-based vaccine delivery needs to be improved to increase the potency of anti-tumor immune response.

**WHAT THIS STUDY ADDS:** NOAH is a novel pipeline for the identification and selection of neoantigens that combines binding affinity and structural features of the peptide/MHC-I interaction. Preclinical studies show highly immunogenic vaccine platform based on HIV-1 Gag based VLPs (neoVLPs) generates antitumor-specific immune responses, delaying tumor growth.

**HOW THIS STUDY MIGHT AFFECT RESEARCH, PRACTICE OR POLICY:** The combination of NOAH and neoVLP platform represents an alternative to current therapeutic vaccine approaches and a promising candidate for future personalized immunotherapy.

## BACKGROUND

Anti-cancer immunotherapies aim to initiate, amplify and expand anti-tumor immune responses (1). Novel therapies that generate *de novo* responses or expand pre-existing neoantigen-specific T cells, with potential to target cancer cells, have proven clinical efficacy in a variety of malignant tumors (2–8). Neoantigens are tumor-specific antigens (TSAs) that derive from single-nucleotide variants (SNVs), altered gene expression (including alternative splicing) or insertions and deletions that lead to frameshifts (9–11). Personalized neoantigen vaccines, which display a limited repertoire of neoepitopes, represent a promising new class of cancer immunotherapy (11–13). Neoantigens are specific to each patient’s tumor and are absent in normal tissues, preventing “off-target” damage (14). Moreover, neoantigen-targeted immune responses bypass central and peripheral tolerance (15).

The identification and selection of neoantigens are critical steps for antitumor vaccine development (15). The field has made significant advancements with the development of next generation sequencing technologies and bioinformatic tools that allow an in-depth analysis of the cancer genome (16). Although mutations play a pivotal role in neoantigen generation, several additional factors are also involved: (i) mRNA expression and its translation into protein, (ii) protein processing, (iii) peptide binding to the MHC and (iv) T-cell receptor (TCR) recognition of the peptide-MHC complex (8,15). Despite each of these events being key, current neoantigen identification strategies have mainly focused on predicting peptide binding to MHC molecules (17), using tools such as NetMHC, NetMHCpan or MHCflurry (18–22). Therefore, further investigation to improve neoantigen identification and selection algorithms is ongoing, including the Tumor Neoantigen Selection Alliance (TESLA) or the NEOantigen Feature toolbOX (NeoFox) (24).

Besides the accurate identification of neoantigens, the success of cancer vaccines also depends on how these neoantigens are formulated and presented to the immune system. Several types of cancer vaccines have reached clinical trials: (i) cell-based vaccines, often prepared as autologous dendritic cells (DCs) pulsed with whole tumor cells, proteins or neoantigens (25–27); (ii) peptide-based vaccines, which induce a robust immune response against the specific tumor antigen-derived peptides (9); (iii) viral vector-based vaccines, such as adenoviruses (28,29); and (iv) nucleic acid-based vaccines, mainly DNA vaccines or the recently developed mRNA technology (7). Remarkably, the combination of different vaccine platforms with immune checkpoint inhibitors, has demonstrated promising results in a phase I clinical trial (30), suggesting that the future of immunotherapies involves the integration of different approaches.

Human Immunodeficiency Virus (HIV-1) Gag-based VLPs constitute a highly suitable vaccine platform to accommodate neoantigens with the aim of generating strong specific T-cell responses with potent antitumor activity. VLPs are complex lipoprotein structures analogous to the corresponding native viruses, but lacking infectivity due to the absence of the viral genome (31,32). HIV-1 Gag-based VLPs are nanoparticles wrapped by a lipid bilayer, similar to retroviruses, that can be generated solely by the expression and subsequent oligomerization of the structural Gag protein monomer (33,34). HIV-1 Gag-based VLPs elicit both humoral and cellular immune responses, exhibit safety, are highly immunogenic and can be produced and purified by standard techniques (34,35). Our research group enhanced the immunogenicity of these HIV-1 Gag-based VLPs (35), which could be further adapted to incorporate specific tumor neoantigens. Therefore, HIV-1 Gag-based VLPs represent an excellent vaccine platform adaptable to mRNA manufacture for the development of personalized cancer vaccines.

Here, we have developed a novel personalized cancer vaccine strategy based on HIV-1 Gag-based VLPs. For that, we used the B16-F10 murine melanoma model to evaluate its efficacy. HIV-1 Gag-based VLPs were engineered to express a collection of neoepitopes that were identified using a novel pipeline including consensus between our novel prediction tool NOAH and existing state of the art software. In contrast to other bioinformatic pipelines, NOAH is not trained on affinity data, which is often associated with high uncertainty, but based on structural features of known peptide/MHC-I interaction. Our results show that vaccinated mice mounted potent neoantigen-specific cellular responses, which were capable of delaying tumor development following inoculation with syngeneic B16-F10 tumour cells.

## METHODS

### Whole exome and RNA sequencing

DNA whole exome libraries of B16-F10 cell line and C57BL/6JOlaHsd germline sample were prepared with Agilent Mouse All Exon kit (Agilent) following manufacturer’s instructions. For RNA sequencing, a total of 1 μg of RNA from the B16-F10 cell line (RIN > 7 and rRNA ratio > 1) was used. RNA library was prepared using the TruSeq Stranded Total RNA Library Prep Gold (Ribozero) kit (Illumina) following manufacturer’s instructions. DNA libraries’ quality control was assessed with Bioanalyzer 2100 (Agilent), quantified by qPCR, normalized and multiplexed into a balanced pool. DNA- and RNA-derived libraries were sequenced on an Illumina NovaSeq600 platform (2x150 paired-end chemistry). Sequencing output of whole exome sequencing (WES) and RNA sequencing (RNAseq) per library yielded 18 Gb (>500X) and 200M reads, respectively.

### In silico neoantigen selection

A pipeline for neoantigen prediction was developed integrating several filters (Figure 1). After an initial variant calling, peptides were ranked by the NOAH algorithm. Briefly (please refer to the results section for more details), NOAH uses a peptide-MHC position specific propensity matrix to rank the peptides, thus inspecting the complementarity between the peptide and the MHC receptor at each amino acid position. Next, ranked peptides by NOAH were crossed with two additional widely used prediction methods, NetMHCpan4 (20) and MHCflurry (22), aiming for consensus. Finally, additional filters were applied: i) having an expression of more than 5 RNA reads, and ii) having a clonality value > 0.2. (variant allele frequency, thus implying that 0.4 of the cells had the variant). NOAH is available for download at https://github.com/BSC-CNS-EAPM/Neoantigens-NOAH.

**Figure 1.**
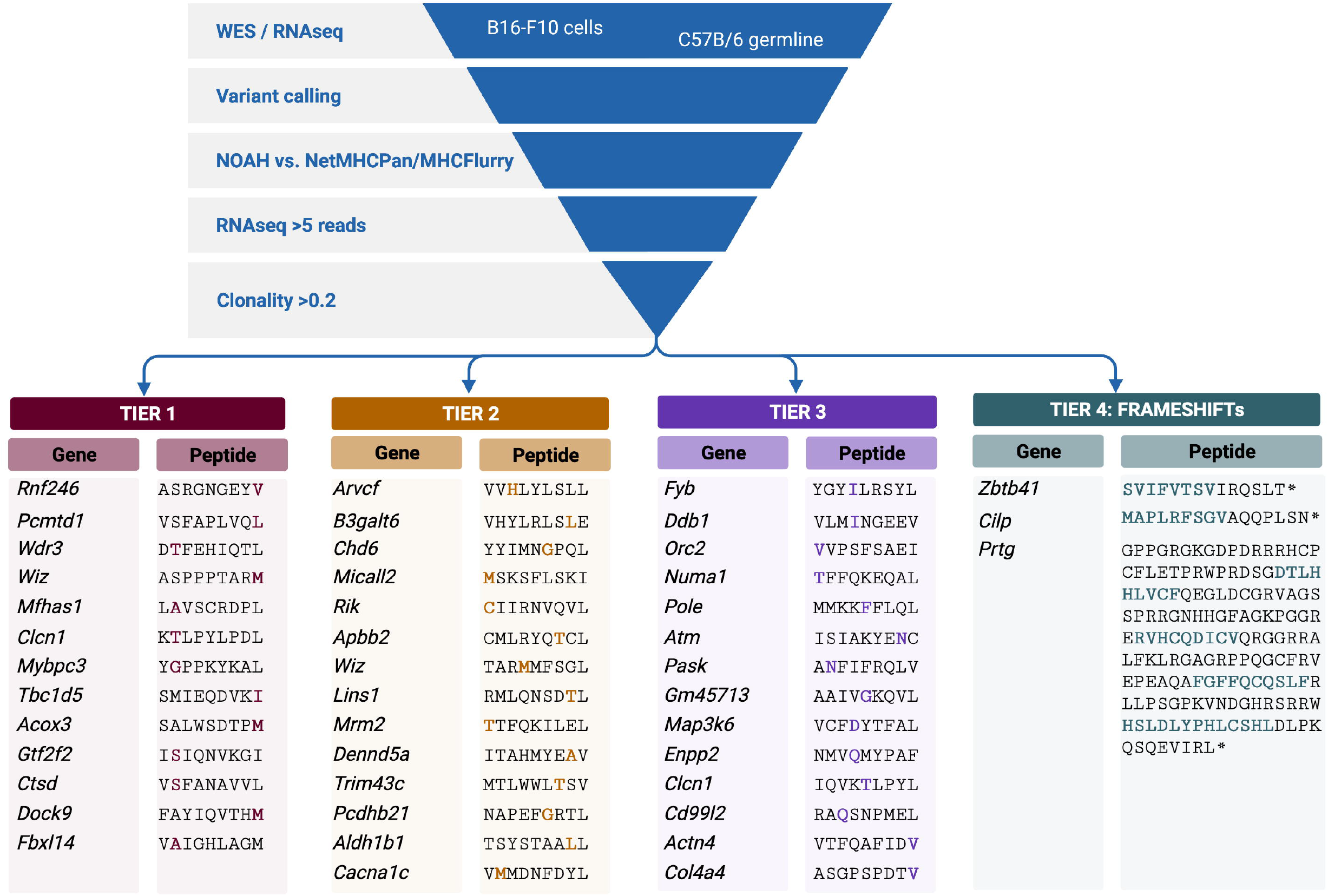
Scheme of the neoantigen selection strategy. Identified somatic mutations were filtered by structural features (NOAH), RNA expression, clonality and matched with NetMHCpan or MHCflurry. Neoantigens tiered according to structural features are shown with the mutation present in B16-F10 cells highlighted in bold. Neoepitopes identified by NOAH in frameshifts are highlighted in bold.

### Plasmids

NeoVLP fusion protein monomers were generated by concatenating from N- to C-term the Flag TAG and the selected neoantigens or frameshifts by an AAA spacer (36), followed by the transmembrane domain of mouse CD44 and by the full sequence of HIV-1 subtype B GAG_HXB2_ (Figure 2A). In the naked-VLP, which acted as a vehicle control, the Flag TAG was directly fused to the murine CD44 transmembrane domain and HIV-1 subtype B GAG_HXB2_ (Figure 2A). All coding sequences were codon optimized and synthetized by GeneArt (Invitrogen), and cloned into pcDNA3.4 (Thermo Fisher). Endotoxin-free plasmids were purified using the ZymoPURE II Plasmid Maxiprep Kit (Zymo).

**Figure 2.**
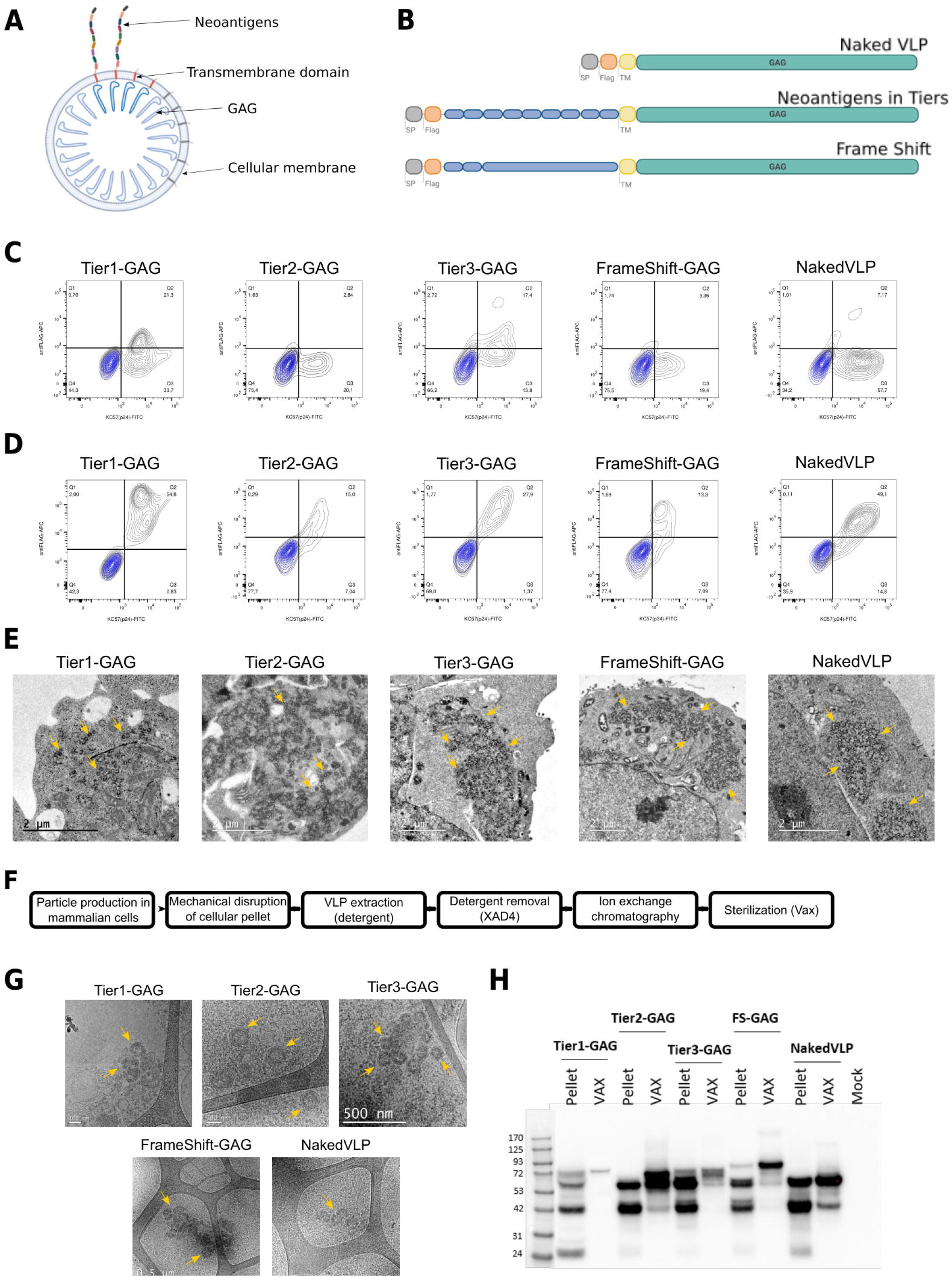
Vaccine platform development based on HIV-1 Virus-Like Particles. **(A)** Scheme of the linear polyprotein that generates the neoVLP. Signal peptide (SP) in light grey, Flag TAG in orange, neoantigens in blue, CD44 transmembrane domain © in yellow and HIV-1 Gag in green. **(B)** Drawing of a neoVLP displaying its components. **(C)** Representative flow cytometry contour plots analyzing the expression of neoVLP fusion proteins in transiently transfected Expi293F cells. Identification of Flag TAG at the surface of the cells and p24-Gag. Mock-transfected Expi293F cells were overlaid in each panel, in blue, for comparison purposes. **(D)** Representative flow cytometry contour plots analyzing the expression of neoVLP fusion proteins in transiently transfected Expi293F cells. Identification of total Flag TAG and p24-Gag. Mock-transfected Expi293F cells were overlaid in each panel, in blue, for comparison purpos© **(E)** TEM images of Expi293F cells producing neoVLP particles. **(F)** Workflow of neoVLP purification. **(G)** Cryo-TEM images of extracted (XAD4) neoVLPs. **(H)** Western blot image evaluating cell lysates (Pellet) and purified neoVLPs (VAX) from each type of VLP.

### Vaccine production and purification

NeoVLPs were produced by transient transfection using Expi293F cells and the Expifectamine293 Transfection Kit following manufacturer’s instructions (Thermo Fisher). Cell cultures were harvested 48h post-transfection. Intracellular neoVLPs were extracted from cell pellets following a previously described protocol (37). Extracted neoVLPs were recovered and loaded on a SepFastDUO5000Q column (BioToolomics). Column flow through was recovered, concentrated by ultrafiltration, filtered at 0.45 μm, and stored at -80°C until use.

### Transmission electron microscopy (TEM)

VLP-producing cells were fixed with 2.5% glutaraldehyde in PBS for 2 hours at 4°C, post-fixed with 1% osmium tetroxide with 0.8% potassium ferrocyanide for 2 hours, and dehydrated in increasing concentrations of ethanol. Then, cell pellets were embedded in EPON resin and polymerized at 60°C for 48 hours. Sections of 70 nm in thickness were obtained with a Leica EM UC6 microtome (Wetzlar), and stained with 2% uranyl acetate and Reynold’s solution (0.2% sodium citrate and 0.2% lead nitrate). Sections were analyzed using a JEM-1400 transmission electron microscope (JEOL) and imaged with an Orius SC1000 CCD Camera (Gatan).

### Cryogenic electron microscopy (Cryo-EM)

VLP morphology was assessed by cryo-EM. Extracted VLPs were deposited on a carbon-coated copper grid and prepared using an EM GP workstation (Leica). Vitrified VLPs were prepared on a Lacey Carbon TEM grid (copper, 400 mesh) and immediately plunge into liquid ethane. The grids were viewed on a JEOL 2011 transmission electron microscope operating at an accelerating voltage of 200 kV. Electron micrographs (Gatan US4000 CCD camera) were recorded with the Digital Micrograph software package (Gatan)

### Flow cytometry

VLP-producing Expi293F cells were extracellularly and intracellularly stained with Allophycocyanin (APC) anti-Flag (DYKDDDDK) tag antibody (1:500) (Biolegend) and intracellularly stained with the Fluorescein isothiocyanate-labeled KC57 (anti-HIV-1 p24) antibody (1:200) (Beckman Coulter) or a mixture of both antibodies. Cells were fixed and permeabilised using the FIX&PERM kit (Invitrogen). Cells were acquired using a BD FACSCelesta Flow Cytometer and data analysis was performed using the Flow-Jo v10.6.2 software (Tree Star Inc.).

### Western blot

Proteins in VLP containing samples were separated by SDS-PAGE using 4-12% Bis-Tris Nu-PAGE gels (Invitrogen) and electro-transferred to a PVDF membrane using the Trans-Blot Turbo Transfer Pack (BioRad). Membranes were blocked (1xPBS pH 7.4, 0.05% Tween20, 5% non-fat skim milk) and subsequently incubated with a rabbit anti-HIV-1 p55+p24+p17 antibody (Abcam, 1:2000) overnight at 4°C. After washing, the membranes were incubated with Peroxidase AffiniPure Goat Anti-Rabbit IgG (H+L) antibody (Jackson ImmunoResearch, 1:10000) for 1h at room temperature (RT), washed and developed using the SuperSignal West Pico PLUS Chemiluminescence Substrate (Thermo Scientific), and images were obtained using a Chemidoc™MP Imaging System (BioRad).

### VLP and total protein quantification

Purified VLPs were quantified either by p24 ELISA (Innotest HIV antigen mAb, Fujirebio) following manufacturer’s instructions or by western blot. For western blot quantification, recombinant Gag protein (35) was used as standard. The standard curve started at 125 ng with 1:2 dilutions until 7.8 ng. Samples were treated as described above. Samples were denatured at 95°C for 5 mins, and proteins were separated by SDS-PAGE. After blocking, membranes were incubated with primary antibody anti-HIV-1 p24 antibody (Abcam, 1:2000) and secondary antibody Peroxidase AffiniPure Donkey anti-Mouse IgG (H+L, Jackson ImmunoResearch, 1:10000).

The total protein content in the sample was assessed by Bicinchoninic Acid (BCA) Protein Assay (ThermoFisher Scientific).

### In vivo experiments

Five-week-old male and female C57BL/6, substrain C57BL/6JOlaHsd, mice were purchased from Envigo. All experimental procedures were performed by trained researchers and approved by the competent authorities (Generalitat de Catalunya, Authorisation ID 9943). All experimental procedures were conducted in accordance with the Spanish laws and the Institutional Animal Care and Ethics Committee of the Comparative Medicine and Bioimage Centre of Catalonia (CMCiB), and following the 3Rs principles. Mice immunization was performed in groups of ten or eight animals. Males and females were equally represented in each group. Mice were firstly immunized with plasmids coding for VLPs (intramuscular electroporation with 20 μg of naked DNA) or with purified VLPs (at the hock, using 100 ng of p24-Gag). Three weeks later, a second dose of vaccine was administered following the same procedure. Blood samples were taken 24 hours before each immunization and tumor cells inoculation. Two weeks after the second immunization, mice were euthanized and a sample of whole blood and the spleen were collected. After blood coagulation (4 hours at RT), serum was collected by centrifugation (10 minutes at 4000xg). Spleens were mechanically disrupted using a 70 μm cell strainer (DDBiolab), and the splenocytes were cryopreserved in FBS containing 10% of dimethyl sulfoxide (Merck).

Two weeks after the second immunization, immunized and control mice were inoculated subcutaneously at the right flank with 10^5^ B16F10 cells (ATCC; CRL-6475) in 100 μL of sterile 1xPBS with 2 mM EDTA. Tumor growth was measured with a caliper every two days and tumor volume (V) was estimated using the formula: *V* = (*length* × *width*^2^) × 0.5, in which length represents the largest tumor diameter and width represents the perpendicular tumor diameter. Humane endpoint was considered when tumor volume was 1 cm^3^ or over. At endpoint, blood samples and spleens were collected and processed, as previously described, for *ex vivo* immune responses analysis.

### Quantification of anti-HIV-1 Gag antibodies by ELISA

The concentration of anti-HIV_Gag_ antibodies in sera of vaccinated mice was determined by ELISA. Nunc MaxiSorp 96-well plates (ThermoFisher Scientific) were coated with 50 ng of recombinant Gag/well (35) in 1xPBS (Gibco) and incubated overnight at 4°C. Coated plates were blocked (1xPBS, 1% of bovine serum albumin (BSA, Miltenyi biotech) and 0,05% Tween20 (Sigma) for 2 hours at RT. Diluted sera (1:100 or 1:1000) from vaccinated mice were loaded onto the plates, incubated overnight at 4°C, washed and incubated with Donkey anti-mouse IgG Fc antibody (Jackson ImmunoResearch, 1:10000) for one hour at RT. Plates were developed using O-phenylenediamine dihydrochloride (OPD, Sigma) and analyzed at 492 nm with a noise correction at 620 nm. As standard reference, anti-HIV-1 p24 antibody (Abcam) was used starting at 333 ng/mL and serially diluted 1:3 down to 0.46 ng/mL.

### Quantification of anti-host cell proteins by Flow cytometry

The humoral response generated against human Expi293F proteins was determined by flow cytometry. Expi293F cells were incubated with mouse serum samples (1:1000) for 30 minutes at RT. After washing, cells were incubated with an AlexaFluor647 goat anti-mouse IgG Fc at a 1:500 dilution (Jackson ImmunoResearch) for 15 minutes at RT. Cells were acquired using a BD FACSCelesta Flow Cytometer and data analysis was performed using the Flow-Jo v10.6.2 software (Tree Star Inc.).

### Quantification of T cell responses by IFNγ ELISpot

Multiscreen ELISpot white plates (Millipore) were coated overnight at 4°C with the anti-mouse IFNγ AN18 antibody (Biolegend) at 2 μg/mL. The following day, plates were washed with sterile PBS containing 1% FBS and blocked with 100 μL of RPMI 1640 medium supplemented with 10% FBS (R10) for 1h at 37°C. After blocking, synthetic peptides (individual neoantigens for Tier1, Tier2 and Tier3; and overlapping peptides for Tier4 (Figure 1) were added at a concentration of 14 μg/mL per peptide, either in individual preparations or in peptide pools. Finally, 4 x 10^5^ splenocytes were added per well and cells were cultured overnight at 37°C. The next day, plates were washed and the biotinylated anti-mouse IFNγ monoclonal antibody R4-6A2 (Biolegend, 1:2000) was added and incubated for 1 hour at RT, followed by an alkaline phosphatase conjugated streptavidin (Mabtech) incubation under the same conditions. IFNγ-specific spots were developed by addition of AP Conjugate substrate Kit (BioRad) and the reaction was stopped by aspiration and incubation for 10 min with 1xPBS (Gibson), 0.05% Tween-20 (Sigma). Concanavalin A (Merck), at 7 μg/mL, was used as a positive control and R10 alone as negative control. Spots were counted using an ELISpot reader S6 Macro M2 (ImmunoSpot, CTL).

### Statistical analysis

Specific CTL responses against individual neoantigen peptides in ELISpot assays were analyzed using Mann-Whitney U test. Multiple comparisons were adjusted by FDR method. Time to sacrifice in each condition were compared by Kaplan-Meier curves and log-rank test.

### Data availability statement

The rest of the data generated in this study are available upon request from the corresponding author, unless stated differently in Materials and Methods particular section.

## RESULTS

### Identification of nonsynonymous mutations and frameshifts in B16-F10 mouse melanoma cell line

To improve the currently available neoantigen selection tools, we set out a novel pipeline that takes into account structural information of the peptide to predict MHC binding. NOAH works under the assumption that binding strength relies on: (i) each position of the peptide; and (ii) the MHC residues that are in contact with each amino acid in the peptide. Thus, NOAH factorizes the peptides into individual (local) positions and builds a position-specific weight matrix (PSMW) mixing validated binding data, whether the peptide binds or not, with structural data from all the reported crystal structures that showed a similar physicochemical space. The final score produced by NOAH is the addition of all local contributions, one per each amino acid in the peptide. Noticeably, this score is not a measure of the binding strength (IC_50_ or percentile rank compared to random peptides), unlike other MHC-binding predictors, but it represents the likeliness of the peptide to properly fit and bind to the MHC. This assumption allows the combination of binding data from different alleles, having similar local environment, and confers a pan-allele status, allowing to also perform *de novo* predictions.

In this study, the B16-F10 melanoma cell line was chosen as a tumor model for the identification of neoantigens. DNA and mRNA were prepared from B16-F10 cells and C57BL/6JOlaHsd healthy tissue and sequenced by WES and RNAseq, followed by variant calling. The mutanome of B16-F10 cells, including SNVs, InDels, and frameshifts, was used to feed NOAH, which gave an output of 51 neoantigen candidates in a ranked manner (SupTable 1). From this candidate list, we selected up to 41 potential neoantigens of 9 amino acids in length (short peptides) from SNVs and three peptides from frameshifts (long peptides), which were grouped into four different tiers (Figure 1). Tier1 emphasized the selection of neoantigens with larger differences on binding affinity between the wild-type and the mutated variant. Neoantigens included in this group presented mutations in MHC anchor residues that are predicted to increase binding to MHC class I molecules. Tier2 grouped neoantigens with high MHC complementarity, as ranked by the consensus approach, bearing mutations that involved a significant change in physicochemical properties (such as polar to aliphatic, negative to positive charge, etc.) for those amino acids that are largely exposed to the solvent and, therefore, are predicted to contact the TCR. Tier3 included peptides that fulfill both binding and expression criteria, but have less drastic changes: with a similar predicted binding to that of the WT and less pronounced changes in a solvent exposed amino acid. Finally, Tier4 included three frameshifts identified by the pipeline and selected for further analysis. The immunogenicity of these selected neoantigens was tested in the context of a novel HIV-1 Gag-based VLP vaccine platform (35) in a syngeneic mouse model.

### Development of HIV-1 Gag-based VLPs carrying neoantigens

Neoantigen-expressing HIV-1 Gag-based VLPs, hereafter called neoVLPs, were engineered to allow a high-density of neoantigens on their surface. Such a high epitope density was obtained by fusing the concatenated neoantigens to the HIV-1 structural protein Gag (35). Since it is estimated that there are around 2500 copies of Gag in one VLP (38), neoVLPs are expected to express the same number of each neoantigen (Figure 2B). NeoVLPs included a signal peptide and a Flag TAG at the N-terminus, followed by the concatenated neoantigen peptides separated by a small spacer sequence (AAA (36) or SSS (39)). This N-terminal sequence was fused to the murine CD44 transmembrane domain followed by the HIV-1 structural protein Gag (Figure 2A). This construct was designed to give rise to a VLP with the N-terminal concatenated neoantigens facing the extracellular space. In this study, three different designs were generated: (i) neoVLPs encoding concatenated neoantigens classified in Tiers 1 to 3 (Tier1-GAG, Tier2-GAG, Tier3-GAG), (ii) a neoVLP encoding the three selected frameshifts in Tier4 (FS-GAG) and (iii) a naked-VLP without neoantigens used as a vehicle control (Figure 2A).

The different fusion constructs were transfected into mammalian Expi293F cells and the expression of the fusion proteins was determined by flow cytometry. The Flag TAG epitope was hardly detected on the cell surface, while both Flag TAG and p24-Gag were readily detected intracellularly (Figure 2C and D), indicating that the fusion proteins were retained inside the cells.

Formation of properly assembled neoVLPs with the expected circular structure in Expi293F cells was demonstrated by transmission electron microscopy (TEM) for each of the fusion proteins tested (Figure 2E). TEM images suggested that the particles budded from the rough endoplasmic reticulum, where the fusion protein was being synthetized and accumulated perinuclearly at the cytoplasm, consistent with a premature association of Gag to intracellular membranes induced by the CD44 membrane spanning domain. No budding events were observed at the plasma membrane, thereby explaining the absence of extracellular Flag TAG staining by flow cytometry.

In order to extract and purify intracellular neoVLPs, transiently transfected Expi293F cells were mechanically disrupted and neoVLPs were extracted by incubation with low detergent concentrations. After detergent removal, neoVLP samples were further purified by multimodal chromatography (strong anion-exchange with a size-exclusion effect) (Figure 2F). Samples from the VLP extracted fraction, prior to the chromatographic step, were imaged by cryo-EM (Figure 2G), displaying the expected morphology for all neoVLPs. From the images, both the lipid bilayer of the enveloped VLP and the electrodense Gag ring inside the generated neoVLPs and naked-VLPs were clearly distinguishable (Figure 2G).

Integrity of the fusion proteins in the cellular lysate and in the final vaccine preparation was evaluated by western blot (Figure 2H). These results confirmed that fusion proteins were produced at the expected molecular weights, even though several bands could be detected, especially in Tier3-GAG lysates, probably due to partial protein processing.

### NeoVLPs induce neoantigen-specific T-cell responses

Next, we tested whether the neoantigens identified *in silico* were immunogenic in the context of natural immunity against B16-F10 tumor cells. To this end, four syngeneic C57BL/6 animals (two males and two females) were inoculated with 10^5^ B16-F10 cells subcutaneously at the right flank (Figure 3A). Mice were euthanized when the tumor volume reached approximately 1 cm^3^, between day 15 and day 20 post-inoculation (Figure 3B). Splenocytes were collected to evaluate neoantigen-specific T-cell responses using IFNγ ELISpot assays. No T-cell responses against any of the selected neoantigens were detected, suggesting that these specificities are not developed during the natural anti-B16-F10 immune responses or are not measurable systemically (Figure 3C).

**Figure 3.**
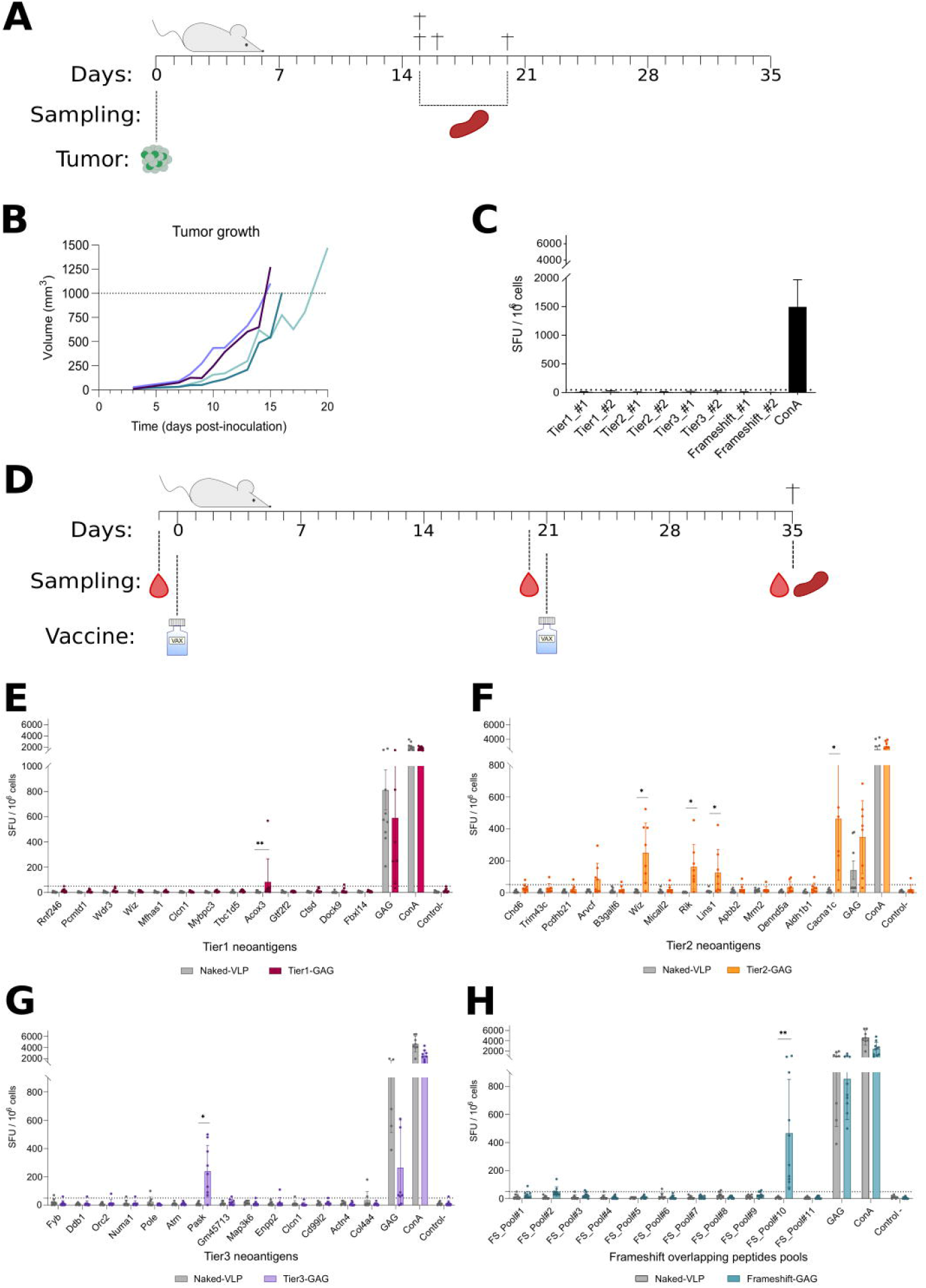
Immunogenicity of selected neoantigens. **(A)** Experimental design for testing natural tumor immunogenicity against selected neoantigens. **(B)** Tumor growth in mice inoculated with 10^5^ B16-F10 cells. Each line represents one animal, two males (dark blue) and two females (light blue) are represented. **(C)** Evaluation of the cellular response against selected neoantigens in mice inoculated with B16-F10 cells. **(D)** Experimental design for testing neoVLP immunogenicity. Blood samples were taken before each vaccination and at endpoint, and spleen was recovered at endpoint. Two vaccines were administered with a three-week interval, and all animals were euthanised two weeks after the second immunization. **(E-H)** Evaluation of cellular responses generated against the selected neoantigens. Tier1-GAG in dark red, Tier2-GAG in yellow, Tier3-GAG in purple, Frameshift-GAG in light blue and naked-VLP in grey.

Then, we tested whether the selected neoantigens, formulated as neoVLPs, could elicit adaptive immune responses by immunization. First, to define the optimal vaccination protocol, C57BL/6 mice were immunized using three different regimens: (i) two doses of naked plasmid DNA coding for VLP protomers (DNA/DNA), (ii) one dose of naked plasmid DNA plus one dose of purified VLPs (DNA/VLP) and, (iii) two doses of purified VLPs (VLP/VLP) (SupFigure 1A). Analysis of the humoral response against HIV-1 Gag protein showed that the DNA/DNA and the DNA/VLP regimes elicited a higher antibody titer, compared to the VLP/VLP regimen (SupFigure 1B). Regarding the generation of cellular immune responses, IFNγ ELISpot analysis against six pools of ten overlapping peptides, in total covering the entire length of the HIV-1 Gag protein, revealed a 10-fold higher CTL response for the DNA/VLP regimen (SupFigure 1C). Therefore, the DNA prime/VLP boost regimen was chosen for immunization in this study. Next, three neoVLPs coding for concatenated neoantigens (Tier1-GAG, Tier2-GAG and Tier3-GAG) and one frameshift (FS-GAG), as well as the naked-VLP, were tested in *in vivo* immunogenicity experiments (Figure 3D). T-cell responses were analyzed by IFNγ ELISpot against individual neoantigen peptides in Tier1-GAG, Tier2-GAG and Tier3-GAG, or against pools of two overlapping peptides for each frameshift in FS-GAG. One single pool of HIV-1 Gag overlapping peptides covering residues 314 to 412 was used in ELISpots as a vaccination positive control for all neoVLPs (Figure 3E-H). T-cell responses were detected against one neoantigen from Tier1-GAG neoVLP, five neoantigens from Tier2-GAG neoVLP, and one from Tier3-GAG neoVLP (Figure 3E-G). Finally, we detected T-cell responses against one out of the three frameshifts tested (Figure 3H), suggesting that peptide length and context might be crucial to induce robust T-cell responses. Therefore, neoantigens classified as Tier2 were the most immunogenic among the selected neoantigens. In addition, immunologically relevant neoantigens were also assessed by IFNγ ELISpot against splenocytes from animals inoculated with B16-F10 cells, which showed an absence of T-cell responses against such neoantigens (SupFigure 2A). All experimental groups generated comparable antibody titers against Gag two weeks after the last vaccination dose (SupFigure 2B), indicating that the differences observed in T-cell responses were not due to variations in vaccine compositions. Accordingly, anti-Expi293F antibodies were also detected in mice immunized with purified VLPs (SupFigure 2C). Taken together, our data suggests that neoVLPs successfully generate *de novo* tumor-specific T-cell immune responses against the selected neoantigens.

### Prophylactic vaccination with neoVLPs delays tumor growth

To determine whether immune responses elicited by neoVLPs were protective against B16-F10-derived tumors, we performed a prophylactic vaccination using Tier2-GAG neoVLPs followed by a B16-F10 tumor challenge assay in syngeneic C57BL/6 mice. Animals were immunized using a DNA/VLP regimen with Tier2-GAG neoVLPs, with or without MPLA as adjuvant. MPLA is a TLR4 agonist inducing Th1 responses (40). A control group immunised with naked-VLP plus MPLA was also included. Two weeks after the vaccine boost (day 35), all mice were inoculated with 10^5^ B16-F10 cells and tumor growth was followed until tumors reached approximately 1 cm^3^ (Figure 4A).

**Figure 4.**
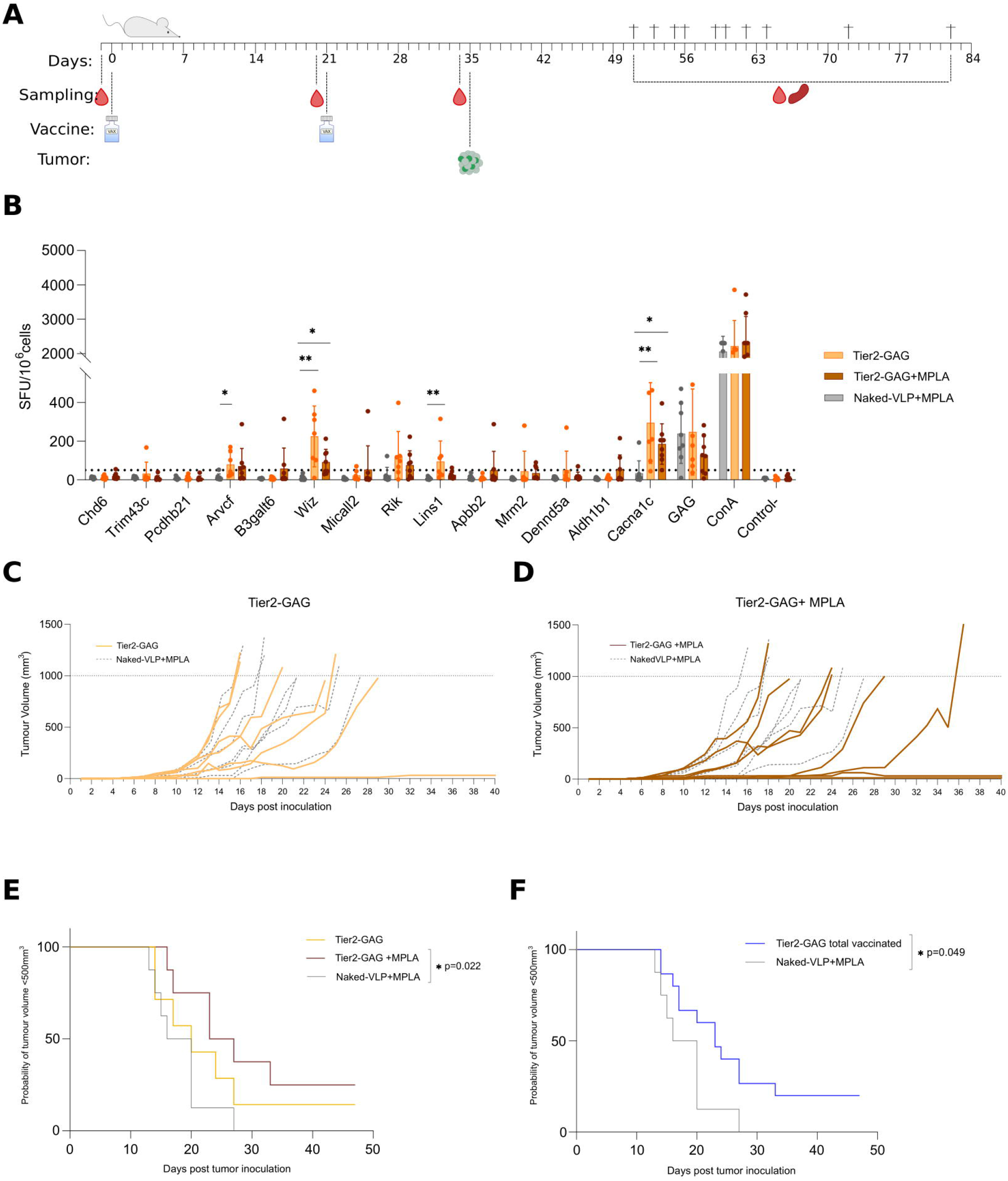
Tumor growth delay and control by neoVLP vaccinated animals. **(A)** Experimental design of a preventive protocol for the evaluation of tumor control. **(B)** Evaluation of cellular responses generated against the selected neoantigens and Gag peptides. **(C)** Tumor growth curves of each animal in the Tier2-GAG group in yellow. Animals vaccinated with naked-VLP are represented by grey dotted lines. **(D)** Tumor growth curves of each animal in the Tier2-GAG+MPLA group in brown. Animals vaccinated with naked-VLP are represented by grey dotted ©es. **(E)** Kaplan-Meier graph representing the time before mice reach a tumor volume equal or over 500 mm^3^. Tier2-GAG in yellow, Tier2-GAG+MPLA in brown and naked-VLP in grey. **(F)** Kaplan-Meier graph representing the time before mice reach a tumor volume equal or over 500mm^3^. Vaccinated with Tier2-GAG in blue (with or without MPLA) and naked-VLP vaccinated mice in grey.

Analysis of the humoral responses showed that all groups generated antibodies against both HIV-1 Gag and Expi293F surface proteins (SupFig 2D and E), whose levels were slightly higher in animals vaccinated with MPLA. In addition, T-cell responses against the previously identified five neoantigens of the Tier 2 group were also detected (Figure 3F and 4B). No effect of MPLA in T-cell responses was observed (Figure 4B). Mice immunized with Tier2 neoVLPs showed a delay in tumor growth compared with control group mice vaccinated with naked-VLPs (Figure 4C-D). In addition, neoVLP-vaccinated animals showed an increased survival rate than control animals (Figure 4E-F). Of note, three animals, one from Tier2-GAG and two from Tier2-GAG+MPLA groups, did not develop any detectable B16-F10-derived tumor (Figure 4C-D). Therefore, our results show that Tier2 neoVLPs promote *de novo* tumor-specific T-cell responses that are capable of generating an anti-tumoral response.

## DISCUSSION

Despite the advances in next generation sequencing techniques and the development of new bioinformatic pipelines for the identification of neoantigens expressed by cancer cells, the identification of strongly immunogenic neoantigens that can develop protective T-cell responses remains challenging due to the low accuracy of the current available pipelines (13,41). Among the different antigen processing steps involved in antigen presentation, the binding of peptides to MHC proteins is considered to be a major determinant. The Immune Epitope Database (IEDB) (42) contains significant noise, i.e., annotations of the same peptide with drastic differences in affinity, which could lead to serious inaccuracies in the training of a peptide binding model against IC_50_ values. In this context, we developed NOAH, a pan-allele method based on a PSWM approach. PSWM methods offer several advantages over machine learning (ML) techniques, including that PSWM: i) are linear and offer a biological explanation of their results, such as residue contribution; (ii) can be trained on qualitative classifications diminishing the impact of experimental errors; (iii) have lower computational requirements than ML methods, allowing a faster screening of peptidomes.

Neural network-based predictions trained on both MHC binding and MHC ligand elution data have achieved the best performance so far in peptide immunogenicity predictions, examples of such pipelines are the well-known NetMHCPan-4.0 or MHCFlurry (43). Even though more than half of the positive predictions of these algorithms or combination of them (MHCcombine) matched with actual binding to the corresponding MHC (43,44), this does not necessarily correlate with a higher immunogenicity of the predicted peptide. In addition, the precision of the predictors in identifying naturally processed MHC-binders is suboptimal compared to predicting binding affinity (45). One reason is the lack of databases reporting the relationship between epitope sequences and the associated T-cell immunogenicity. Alternatively, structure-based predictions can provide high-resolution TCR-peptide-MHC structure (45), which allows a better assessment of the interaction with the TCR and, therefore, the immunogenicity of the predicted epitope. Here, we have developed a novel neoantigen selection pipeline which not only takes into account the binding affinity and the complementarity between the peptide and the MHC, but also its interaction with the TCR, by focusing the selection on some specific peptide positions and the physicochemical properties of the variation.

Furthermore, to overcome the low immunogenicity associated with peptide immunization, we have generated a novel HIV-1 Gag-based VLP platform that can accommodate several neoantigens at high density within each particle, with the aim of increasing its immunogenicity (35). The *in vivo* immunogenicity of NOAH-predicted neoantigens was tested using this novel vaccine platform. We classified neoantigens identified *in silico* into three Tiers based on the type of mutation and its location: whether it affects MHC binding or interaction with the TCR, or depending on its similarity to the wild-type sequence. In addition, we included frameshift mutations as a fourth-Tier category. Nonetheless, our results showed that neoantigens classified mainly in Tier2, which contained drastic amino acid changes in a position that is likely to be in contact with the TCR, were able to generate stronger T cell responses after immunization with neoVLPs. These data emphasize that beyond the binding affinity to the MHC-I, the interaction of the MHC-I/neoantigen complex with the TCR is key for neoantigen identification. Frameshift mutations generate a complete change in the amino acid sequence of the affected protein compared to its wild-type counterpart. Consequently, frameshifts are expected to be a reliable source of immunogenic neoantigens (46,47). In this study, we included three frameshifts for *in vivo* experimental validation. Our results confirm frameshift mutations as a good source of immunogenic neoantigens.

Remarkably, T-cell responses against the selected neoantigens were not detected in mice bearing the tumor, suggesting either that they are not the main target of the natural anti-tumor immune response in these animals or that B16-F10 tumor cells are poorly immunogenic. That is consistent with the high aggressiveness displayed by B16-F10 cells in C57BL/6 mice and their limited response to checkpoint inhibitors (29). Notably, Tier2-GAG vaccinated animals showed delayed tumor growth and increased survival. In fact, three out of sixteen animals did not develop the tumor, indicating that Tier2-elicited T-cell responses may be protective. Although the vaccination alone has demonstrated to be insufficient to protect all animals, the generation of novel neoantigen-specific T-cell responses indicate that the protective effect observed with Tier2-GAG VLPs may be enhanced by combining with immune checkpoint inhibitors, as it has been demonstrated in a therapeutic setting (30). However, further work is needed to confirm the efficacy of neoVLPs in combination with other currently available immunotherapies, such as immune checkpoint inhibitors or inflammatory cytokines, such as IL-2.

In summary, our findings provide a promising strategy for the development of personalized cancer vaccines. We have presented an innovative *in silico* neoantigen selection pipeline based on a novel peptide-MHC binding predictor, NOAH, and a consensus approach. In addition, we have adapted our HIV-1 Gag-based VLP vaccine platform for the generation of protective neoantigen-specific cellular immune responses in mice. Overall, these results confirm that neoVLPs are promising candidates for future personalized immunotherapies against cancer.

## Acknowledgements

We would like to thank the staff from the Servei de Microscopia of Universitat Autònoma de Barcelona. We are grateful to CMCiB staff for their excellent technical help with animal care. Finally, we would like to extend our sincere thanks to Dr Francesc Cunyat for reviewing the manuscript and providing valuable feedback.

---

**Supplementary Figure 1. Selecting the vaccination regimen for the highest immune response. (A)** Experimental design for testing neoVLP vaccine regimen. **(B)** Evaluation of the humoral response generated against recombinant Gag at sacrifice. DNA/DNA regimen in purple, DNA/VLP regimen in blue and VLP/VLP regimen in turquoise. **(C)** Evaluation of T cell response against pools of peptides covering the HIV-1 Gag protein. DNA/DNA regimen in purple, DNA/VLP regimen in blue and VLP/VLP regimen in turquoise.

**Supplementary Figure 2. Natural tumor immunogenicity against neoantigens and humoral response against Gag and host cell proteins. (A)** Cellular response generated by natural tumor immunogenicity against selected neoantigens. **(B)** Evaluation of humoral response against HIV-1 Gag over time for all groups vaccinated with neoVLPs. **(C)** Evaluation of the humoral response against Expi293F proteins at each endpoint for all groups vaccinated with neoVLPs. Level of response in vaccinated animals is displayed as coloured dots according to each group. Staining controls are shown as grey dots. **(D)** Evaluation of humoral response against HIV-1 Gag for groups vaccinated with Tier2-GAG (in yellow), Tier2-GAG+MPLA (in brown) and naked-VLP+MPLA. **(E)** Evaluation of the humoral response against host proteins at endpoint for groups vaccinated with Tier2-GAG (in yellow), Tier2-GAG+MPLA (in brown) and naked-VLP+MPLA (in grey).

**Supplementary Table 1. B16-F10 cells mutanome**. Gene, neoantigen and wild-type peptides are indicated. NOHA, MHCflurry and NetMHCpan4 scores are also shown for H2-Db and H2-Kb mouse MHC-I.

## Notes

**Competing interests** The authors declare that the research was conducted in the absence of any commercial or financial relationships that could be construed as a potential conflict of interest. Outside this work BC, JB and JC are founders and shareholders of AlbaJuna Therapeutics, S. L.

**Funding.** This work was supported by the CERCA Program (2017 SGR 252; Generalitat de Catalunya), PID2019-106370RB-I00/AEI/10.13039/501100011033 (Spanish Ministry of Science and Innovation), Grifols S. A. (Spain) and Sorigué (Spain). Funders had no role in the study design, data analysis, decision to publish, or manuscript preparation. AP-G was funded by Secretaria d’Universitats i Recerca of Generalitat de Catalunya and the European Social Fund through the fellowship 2022FI_B00698.

### Competing Interest Statement

The authors declare that the research was conducted in the absence of any commercial or financial relationships that could be construed as a potential conflict of interest. Outside this work BC, JB and JC are founders and shareholders of AlbaJuna Therapeutics, S. L.

