## Supplementary figures and images for "VLP-mediated delivery of structure-selected neoantigens demonstrates immunogenicity and antitumoral activity in mice"

### Supplemental Figure 1

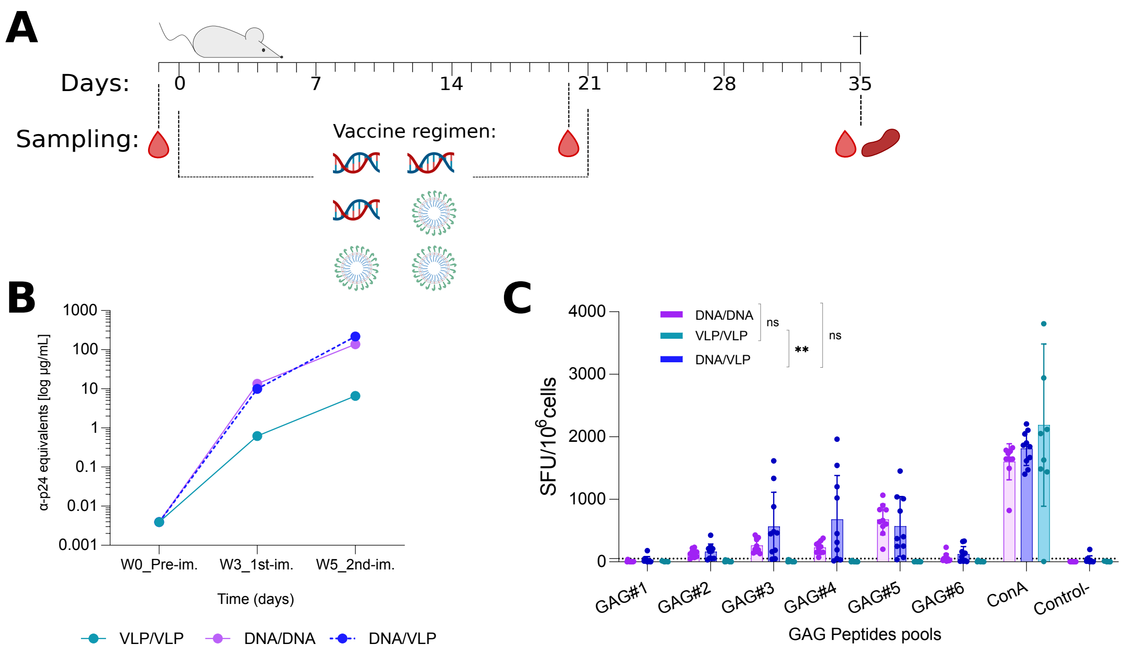

### Supplemental Figure 2

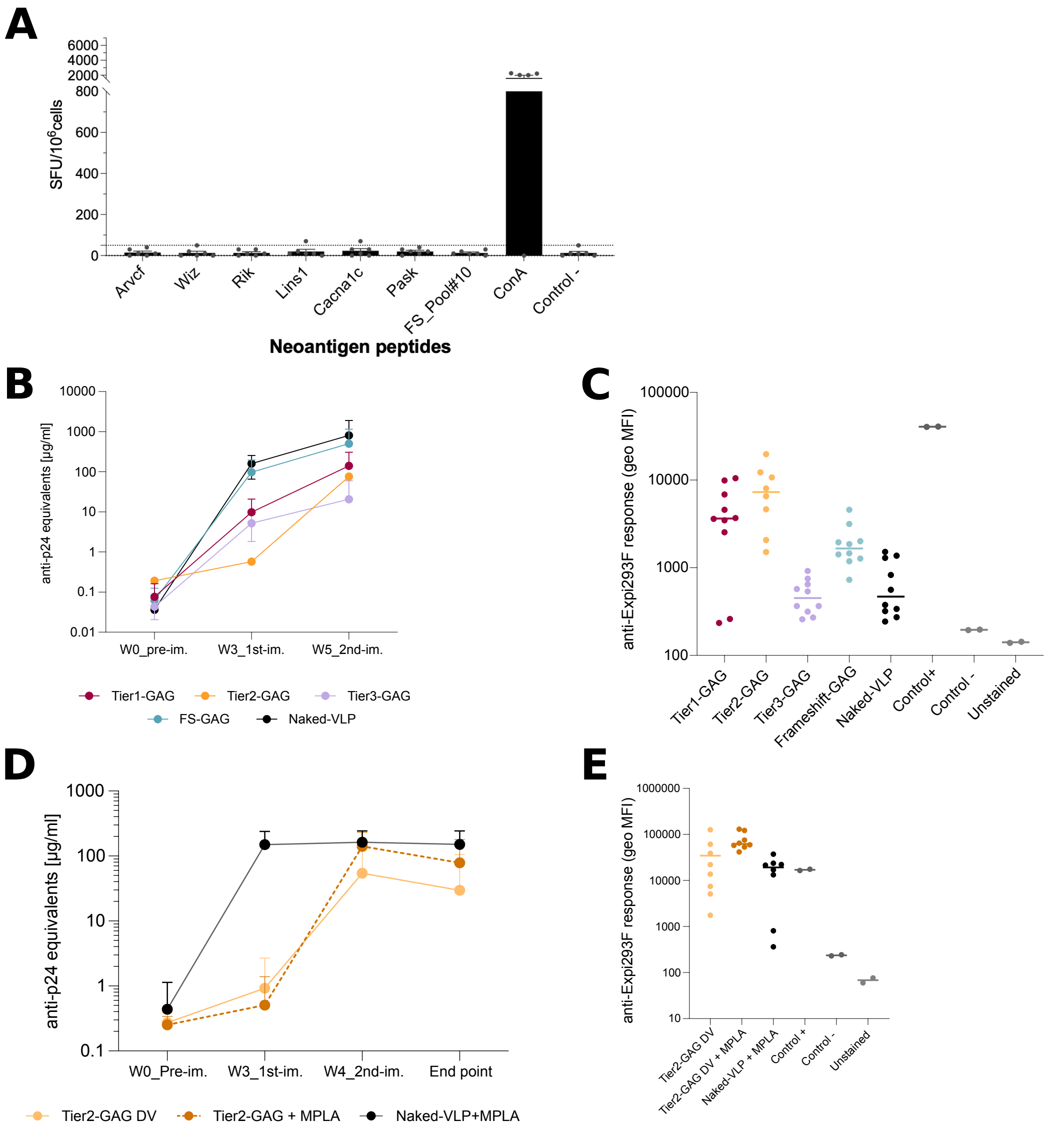
